# The use of cultured human alveolar basal cells to mimic honeycomb formation in idiopathic pulmonary fibrosis

**DOI:** 10.1101/2023.09.19.557680

**Authors:** Sabrina Blumer, Petra Khan, Nataliia Artysh, Linda Plappert, Spasenija Savic, Lars Knudsen, Danny Jonigk, Mark P. Kuehnel, Antje Prasse, Katrin E. Hostettler

## Abstract

Honeycomb cysts (HC) within the alveolar region are distinct histopathological features in the lungs of idiopathic pulmonary fibrosis (IPF) patients. HC are lined with basal cells (BC), or with a bronchiolar-like epithelium composed of basal-, ciliated- and secretory epithelial cells. By using cultured IPF patient-derived alveolar BC, we aimed to establish *in vitro*- and *in vivo* models to mimic HC formation in IPF. In order to do so, we cultured the cells (1) on an air liquid interface (ALI) or (2) in a three dimensional (3D) organoid model *in vitro*, and (3) investigated the cells’ behavior after instillation into bleomycin-challenged mice *in vivo*. Under the here tested *in vitro*- and *in vivo* conditions, alveolar BC differentiate and formed HC-like structures, which closely resemble HC within the IPF lung. These models therefore represent powerful tools to study HC formation, and its potential therapeutic inhibition in IPF.

## Introduction

Idiopathic pulmonary fibrosis (IPF) is characterized by progressive destruction of the lung parenchyma and the respective irreversible loss of lung function (1). Repetitive injuries to the alveolar epithelium and its disturbed regeneration play a critical role in disease development (2). In the healthy lung, the alveoli are lined with alveolar epithelial cells type 1 and 2 (AT1 and 2) that facilitate gas exchange and provide the lung with the four essential surfactant proteins (SP)-A, -B, -C, and -D, of which SP-C is specifically expressed in AT2 cells (3). After mild-to-moderate lung injury, AT2 cells regenerate the alveolar epithelium by their ability to self-renew and to differentiate into AT1 cells (3). However, chronic lung injury in IPF results in a maladaptive repair mechanism, in which the alveolar lung parenchyma is replaced by dense fibrotic tissue and honeycomb cysts (HC). HC are lined with a single-, or stratified layer of KRT (keratin)5+ basal cells (BC) or by a stratified bronchiolar-like epithelium composed of KRT5+ basal-, acetylated tubulin (AcTub)+ ciliated-, and secretory epithelial cells, mainly expressing MUC (mucin) 5B (4–9). Although, the appearance of ectopic airway epithelial cells within the peripheral lung of IPF patients is a well described phenomena, the cells origin and functional role in disease development and progression remains largely unknown. This may partly be explained by a lack of *in vitro* and *in vivo* models that appropriately resemble HC formation in IPF.

We previously reported the fibrosis-enriched outgrowth of cells with distinct morphology from peripheral lung tissue of IPF patients (10). Characterisation of the cells revealed their expression of canonical BC markers and close transcriptomic similarity to BC (6). Under the previously described cell culture conditions, the cells expansion and experimental use proved difficult (6, 10, 11). In this study, we therefore first optimized the culture of alveolar BC and then used the cells to establish *in vitro* and *in vivo* models to mimicking HC formation in IPF.

## Results

### The alveolar lung parenchyma is replaced by dense fibrotic tissue and HC in IPF lung tissue

Compared to the normal alveolar lung parenchyma with some peripheral bronchioles in close proximity to pulmonary arteries in non-fibrotic lung tissue of control patients (n=4), the alveolar lung parenchyma in IPF patients is destroyed and replaced by dense fibrosis with multiple HC (n=4) (Figure 1 A). IPF lung tissue (n=4) displayed a reduced number of SP-C+ AT2 cells when compared to non-fibrotic lung tissue of control patients (n=4) (Figure 1 B, D). Cells positive for the canonical basal cell marker KRT17, KRT5 and tumor protein 63 (p63), or secretory (secretoglobin family 1A member 1 (SCGB1A1)+, MUC5B+)- or ciliated (AcTub+) epithelial cells were detected within distal bronchioles of non-fibrotic controls tissue and within HC in IPF tissue (Figure 1 B). Overall numbers of KRT17-, KRT5-, p63-, SCGB1A1-, MUC5B-, or AcTub-expressing epithelial cells was significantly higher in IPF when compared to control tissue (Figure 1 C, D). KRT14 was only expressed in a fraction of cells within IPF, but not in control lung tissue (Figure 1 C, D). MUC5AC+ cells were absent in the alveolar epithelium of control tissue and only detected in small numbers in IPF lungs (Figure 1 C, D). Matrix metalloprotease (MMP)7, an established biomarker for IPF (12), was only detected in IPF, but not in control lung tissue (Figure 1 C, D).

**Figure 1:**
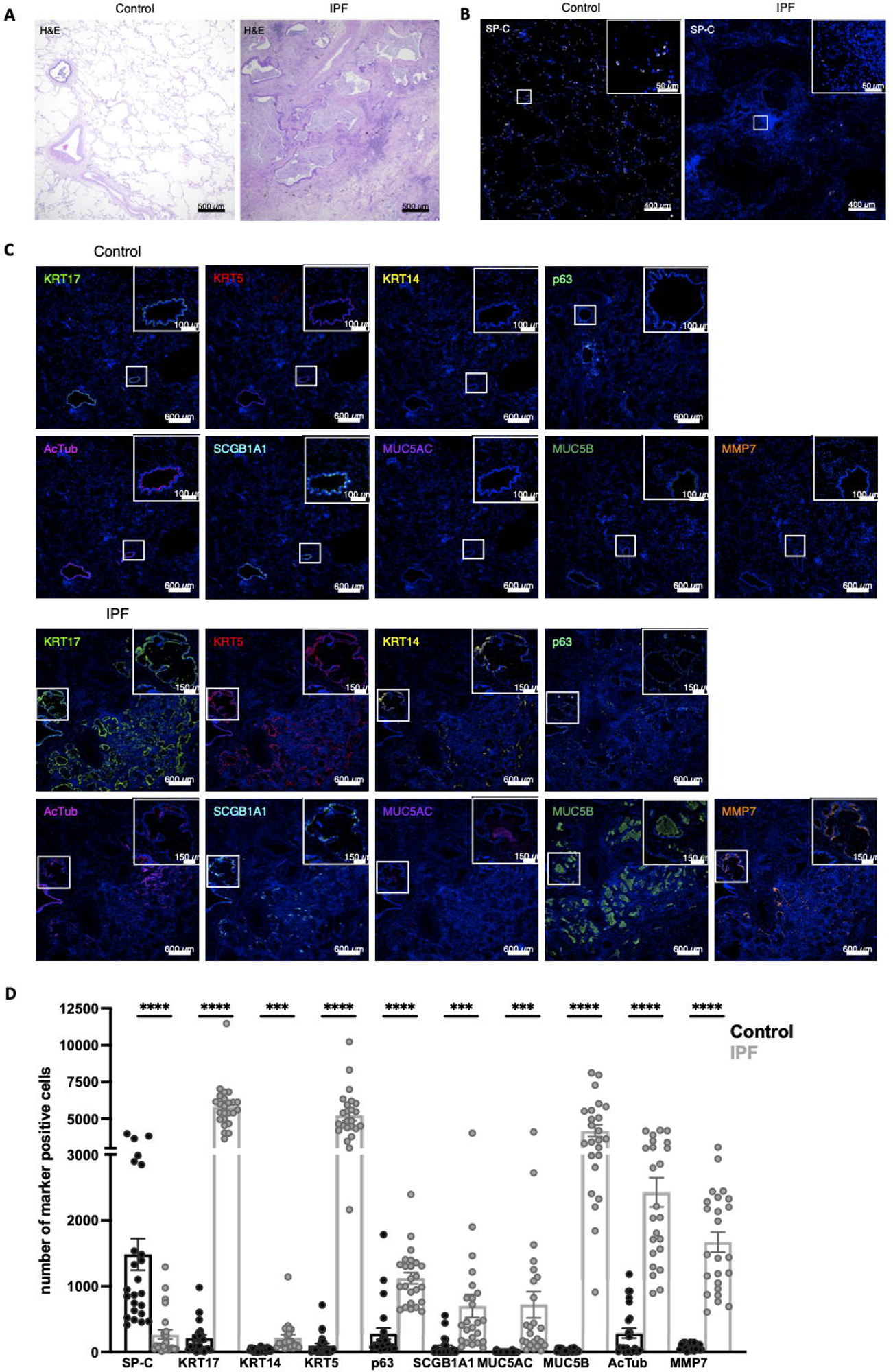
**Decreased number of AT2 cells is accompanied with an accumulation of airway epithelial cells in peripheral IPF lung tissue**. Representative images showing Hematoxylin and Eosin (H&E) stainings in control (n=4) and IPF (n=4) lung tissue (A). Representative IHC/IF images of control (n=4) and IPF (n=4) lung tissue incubated with antibodies detecting SP-C (B), MMP7, or basal (KRT5, KRT17, KRT14, p63)-, ciliated (AcTub)-, or secretory (SCGB1A1, MUC5AC, MUC5B) epithelial cell markers (C). White squares indicate regions imaged at higher magnification shown in the top right corner of the image (B, C). IHC/IF image quantifications of cells expressing specific markers within control (n=4) and IPF (n=4) tissue (D). Dots within the bar charts represent individual data points generated by analysis of six images of control (n=4) and IPF (n=4) lung tissue. Data are expressed as mean ± SEM (unpaired t-test, **** p < 0.0001, *** indicates p < 0.001).

### Cultured alveolar BC express high levels of basal cell markers, and show a robust proliferation and wound healing capacity

Peripheral IPF lung tissue was cut in small pieces and placed in cell culture dishes containing an epithelial cell specific growth medium (Cnt-PR-A) as illustrated in Figure 2 A. Outgrown alveolar BC expressed high levels of TP63 (gene name of the p63 protein), KRT5, KRT14, and KRT17 on the RNA (Figure 2 B) and protein (Figure 2 C) level. RNA of the secretory epithelial cell marker SCGB1A1 or mesenchymal marker CDH2 was either not expressed or expressed at very low levels (Figure 2 B). SCGB1A1 or CDH2 proteins were absent in cultured cells (Figure 2 C).

**Figure 2:**
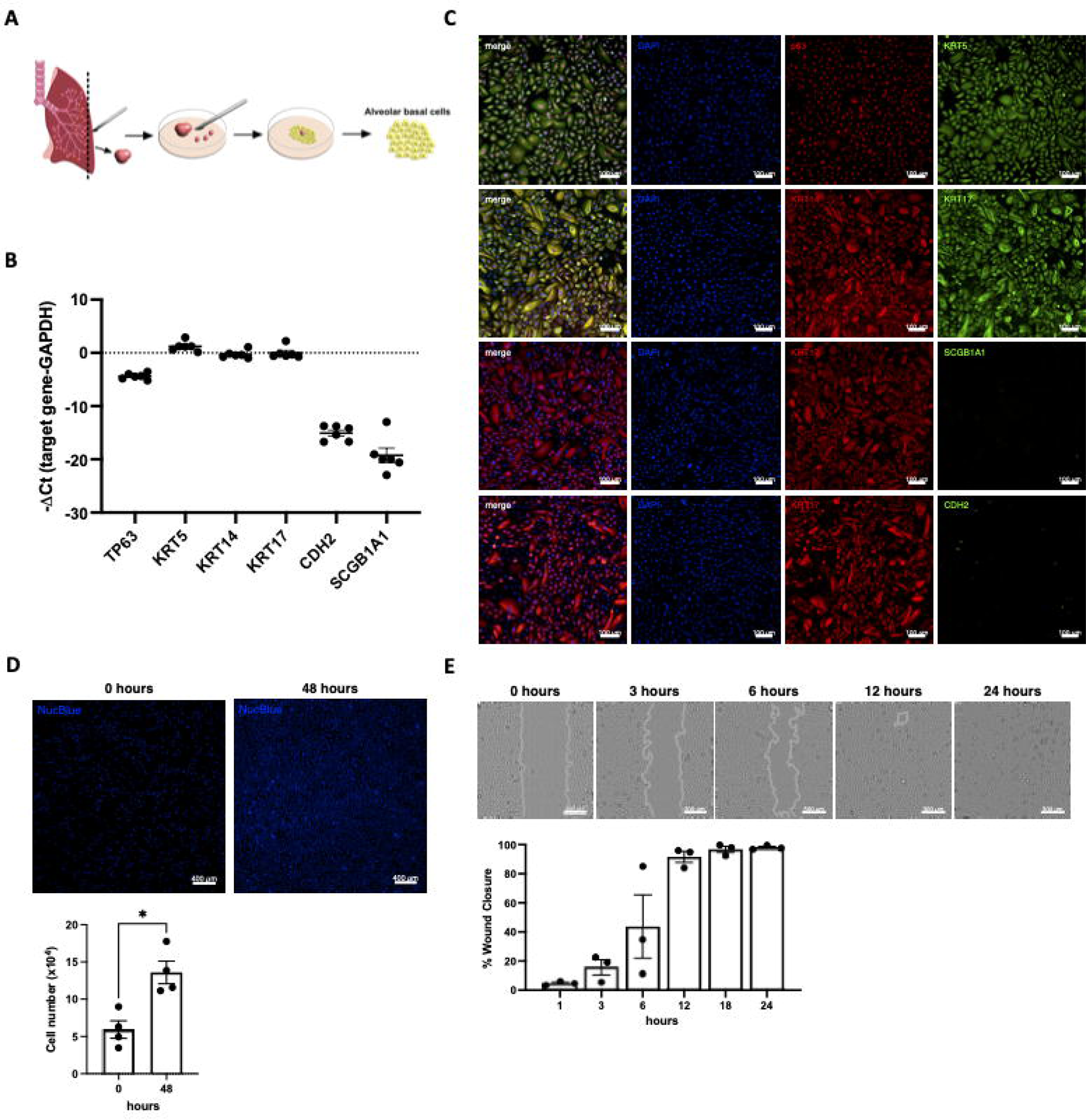
Characteristics of cultured alveolar basal cells. Illustration of alveolar BC culture (A). Experiments presented in this figure were performed in alveolar BC from 3-6 different IPF patients (n-number). RNA expression levels (-ΔCt) of TP63, KRT5, KRT14, KRT17, CDH2 and SCGB1A1 in alveolar BC (n=6). Data are expressed as mean ± SEM (B). Representative single- or merged ICC/IF images of alveolar BC (n=4) incubated with antibodies detecting KRT5, p63, KRT17, KRT14, SCGB1A1, or CDH2. Nuclei were counter-stained with DAPI. Automated cell counts of alveolar BC at 0 and 48 hours (n=4) (D). Alveolar BC wound closure between 0-24 h (n=3) (E). Dots within the bar charts represent datapoints generated for alveolar BC from each patient. Data are expressed as mean ± SEM (paired t-test, * indicates p < 0.05).

Cultured alveolar BC displayed robust proliferation, shown by a significant increase in cell numbers between 0 to 48 hours (Figure 2 D) and wound healing capacity by closing a mechanically set scratch after 12 hours (Figure 2 E).

### Alveolar BC can be propagated for up to six passages and cryopreserved without significant loss of basal cell markers

Confluent alveolar BC were trypsinized and split at a ratio of 1 to 2 for up to six passages or were cryopreserved in cryo-medium. Between passage 0-4, no reduction in the expression of basal cell marker KRT5 and TP63 or that of the canonical epithelial cell marker CDH1 was detected on the RNA (Figure 3 A) or protein level (Figure 3 B). Between passage 5-6 a slight, but insignificant decrease in KRT5 and TP63 expression was observed on the RNA and protein level (Figure 3 A, B). The mesenchymal marker CDH2 was detected at very low levels on the RNA level (Ct values >30) (Figure 3 A) and was absent on the protein level in cells between passage 1-5 (Figure 3 B). Similarly, alveolar BC before (fresh) and after cryopreservation (thawed) did not show any significant changes in KRT5, TP63, CDH1, or CDH2 RNA expression (Figure 3 D).

**Figure 3:**
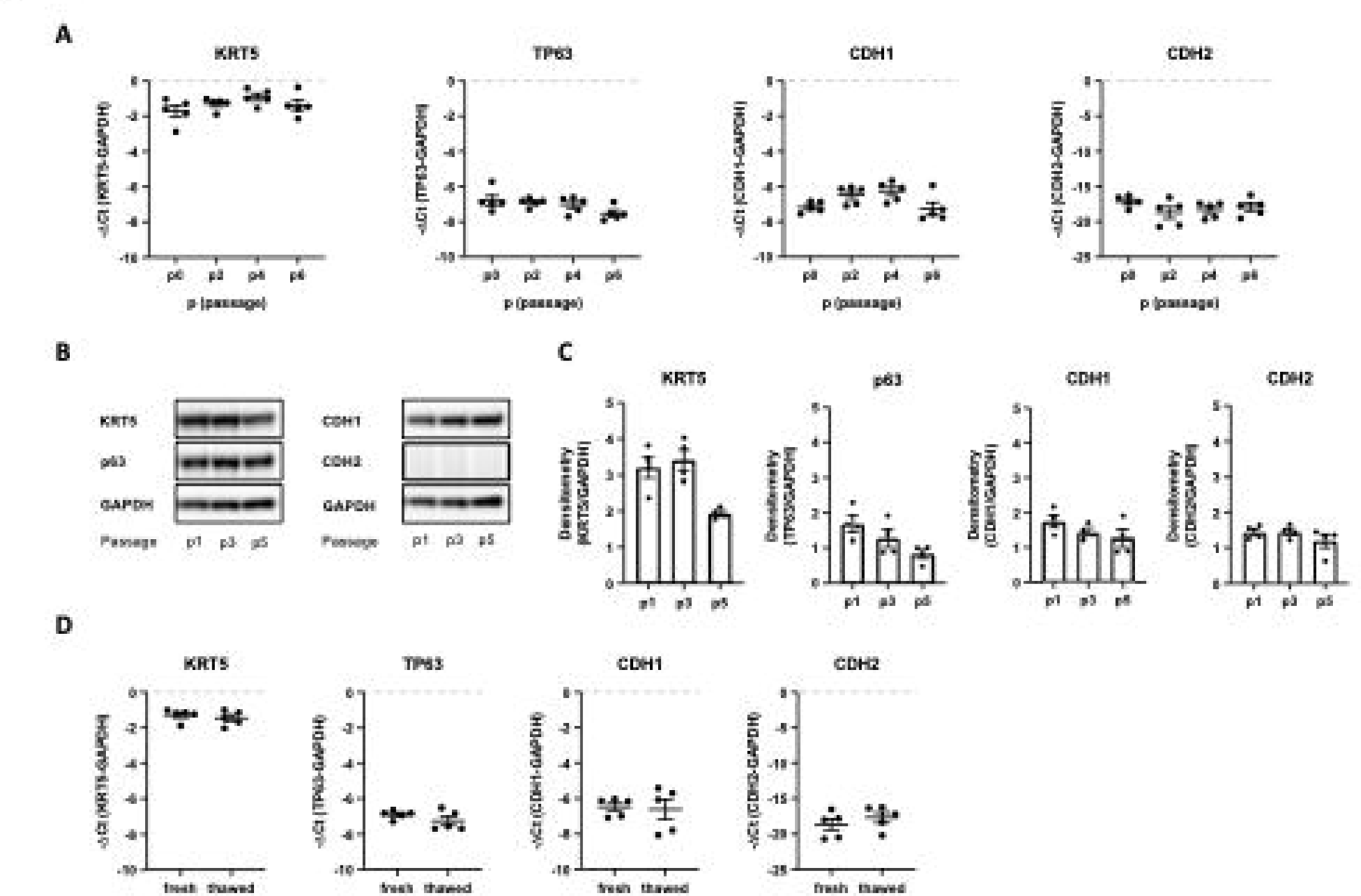
Alveolar basal cell marker expression is stable over several passages and after cryopreservation. Experiments presented in this figure were performed in alveolar BC from 4-5 different IPF patients (n-number). RNA expression levels (-ΔCt) of KRT5, TP63, CDH1 or CDH2 in alveolar BC (n=5) at passage (p) 0, 2, 4, and 6 (A). Representative immunoblots showing the protein expression of KRT5, p63, CDH1, CDH2 or GAPDH in alveolar BC (n=4) at p1, p3, and p5 (B) and their densitometry analysis relative to the house keeping gene GAPDH. Dots within the bar charts represent datapoints generated for alveolar BC from each patient (C). RNA expression levels (-ΔCt) of KRT5, TP63, CDH1 or CDH2 in alveolar BC (n=5), before (fresh) and after (thawed) cryopreservation (D). Data are expressed as mean ± SEM (A, C, D).

### Alveolar BC differentiate towards ciliated- and secretory epithelial cells when cultured on an ALI or in a 3D organoid model

Illustration of mucociliary differentiation of alveolar BC cultured on an ALI platform for 23 days (Figure 4 A). At day 0, RNA expression of the basal cell marker KRT5 is high and that of MMP7, secretory (SCGB1A1, MUC5AC, MUC5B)- or ciliated (FOXJ1) epithelial cell markers are absent or expressed at low levels (Figure 4 B). After 23 days of ALI culture, RNA of MMP7 and all secretory- and ciliated epithelial cell marker are up-regulated and that of KRT5 down-regulated (Figure 4 B). Similarly, SCGB1A1-, MUC5AC-, and AcTub proteins are absent in cells at day 0 and abundantly present in cells at day 23 (Figure 4 C). Immunofluorescence stainings of transwell membrane cryosections show alveolar BC differentiation into a stratified epithelial cell layer with KRT5+/KRT17+/p63+ basal cells at the basal region and secretory (SCGB1A1+, MUC5AC+, MUC5B+, MMP7+)-, and ciliated (AcTub+) epithelial cells at the apical region (Figure 4 D).

**Figure 4:**
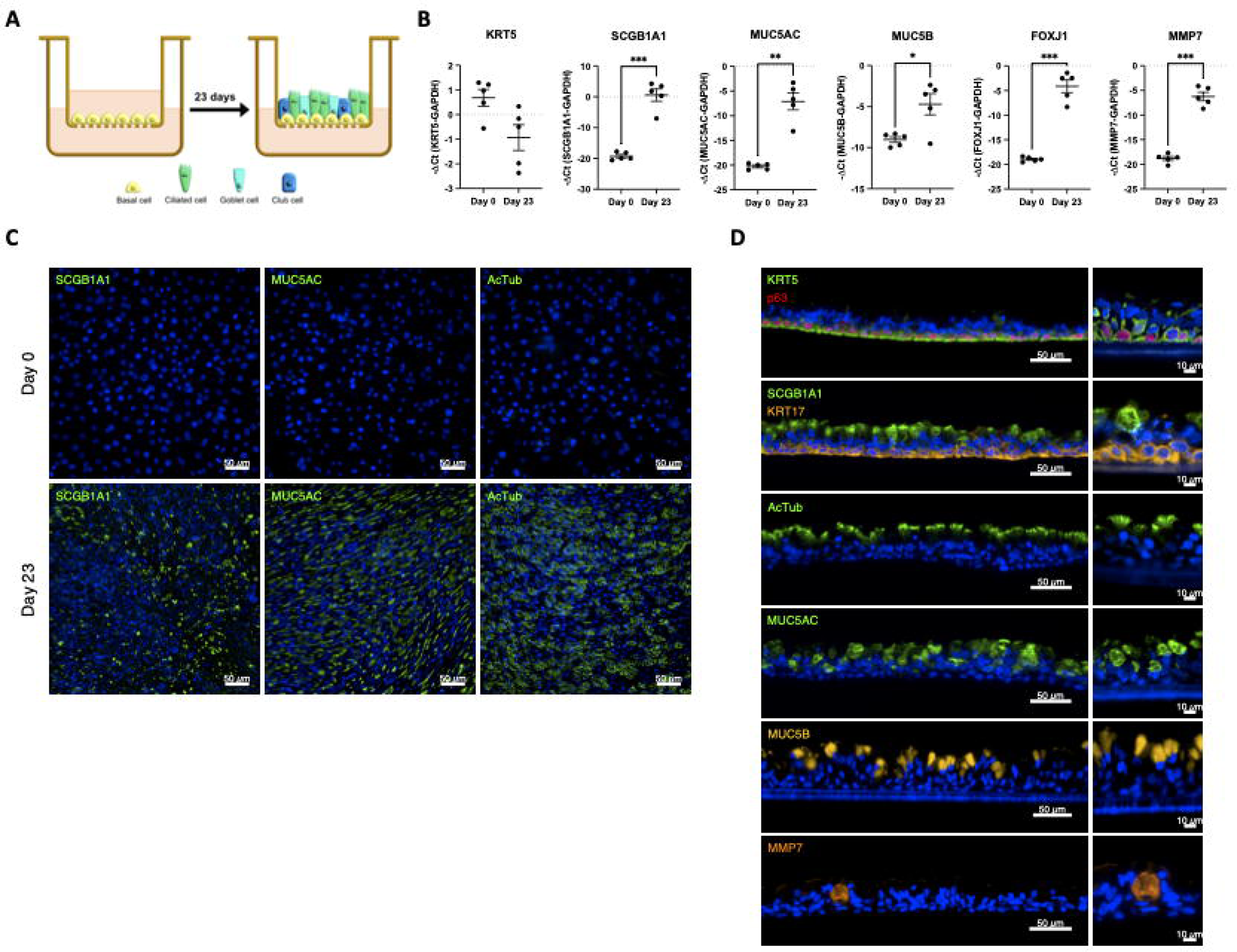
Alveolar basal cells cultured on an ALI differentiate to secretory- and ciliated epithelial cells. Illustration of alveolar BC ALI culture (A). Experiments presented in this figure were performed in alveolar BC from 3-5 different IPF patients (n-number). RNA expression levels (-ΔCt) of KRT5, SCGB1A1, MUC5AC, MUC5B, FOXJ1, or MMP7 in alveolar BC (n=5) cultured on an ALI on day 0 and day 23. Data are expressed as mean ± SEM (paired t-test, *** indicates p < 0.001, ** indicates p < 0.01, * indicates p < 0.05) (B). Representative ICC/IF images of ALI cultured alveolar BC (n=2) incubated with antibodies detecting SCGB1A1, MUC5AC, or AcTub at day 0 and day 23 (C). Representative images of ALI culture cryosections (n=3-4) incubated with antibodies detecting KRT5, KRT17, p63, SCGB1A1, MUC5AC, AcTub, MUC5B or MMP7 at day 23 (D). Nuclei were counter-stained with DAPI (C, D).

Alveolar BC were embedded in Matrigel and cultured for 21 days (Figure 5A). Cells expand in numbers and start to self-assemble into clusters at day 2. Organoids (>50µm diameter) first formed after 7 days and continued to grow over a period of 21 days (Figure 5 B). After 20 days, 43-100 organoids/mm^3^ with a diameter of 50-280µm were formed (Figure 5 C). Cells within organoids expressed KRT17-, KRT5-, TP63-, SCGB1A1-, MUC5AC-, MUC5B-, FOXJ1-, and MMP7 RNA (Figure 5 D). Furthermore, cells showed positive IF signals for KRT17-, KRT5-, p63-, AcTub-, MUC5AC-, MUC5B-, and MMP7 as presented in figure 5 E and quantified in figure 5 F. Polarized lumen were present in approximately 40% of the organoids after 21 days, with BC (KRT5+, KRT17+, p63+) mainly located at the outer layer, and ciliated (AcTub+)-, secretory (SCGB1A1+, MUC5AC+, MUC5B+)-, or MMP7+ epithelial cells at the luminal site of the organoids. Similar to HC within the IPF lung (Figure 1 C), the lumen of the organoids were filled with MUC5B+ or MUC5AC+ mucus (Figure 5 G).

**Figure 5:**
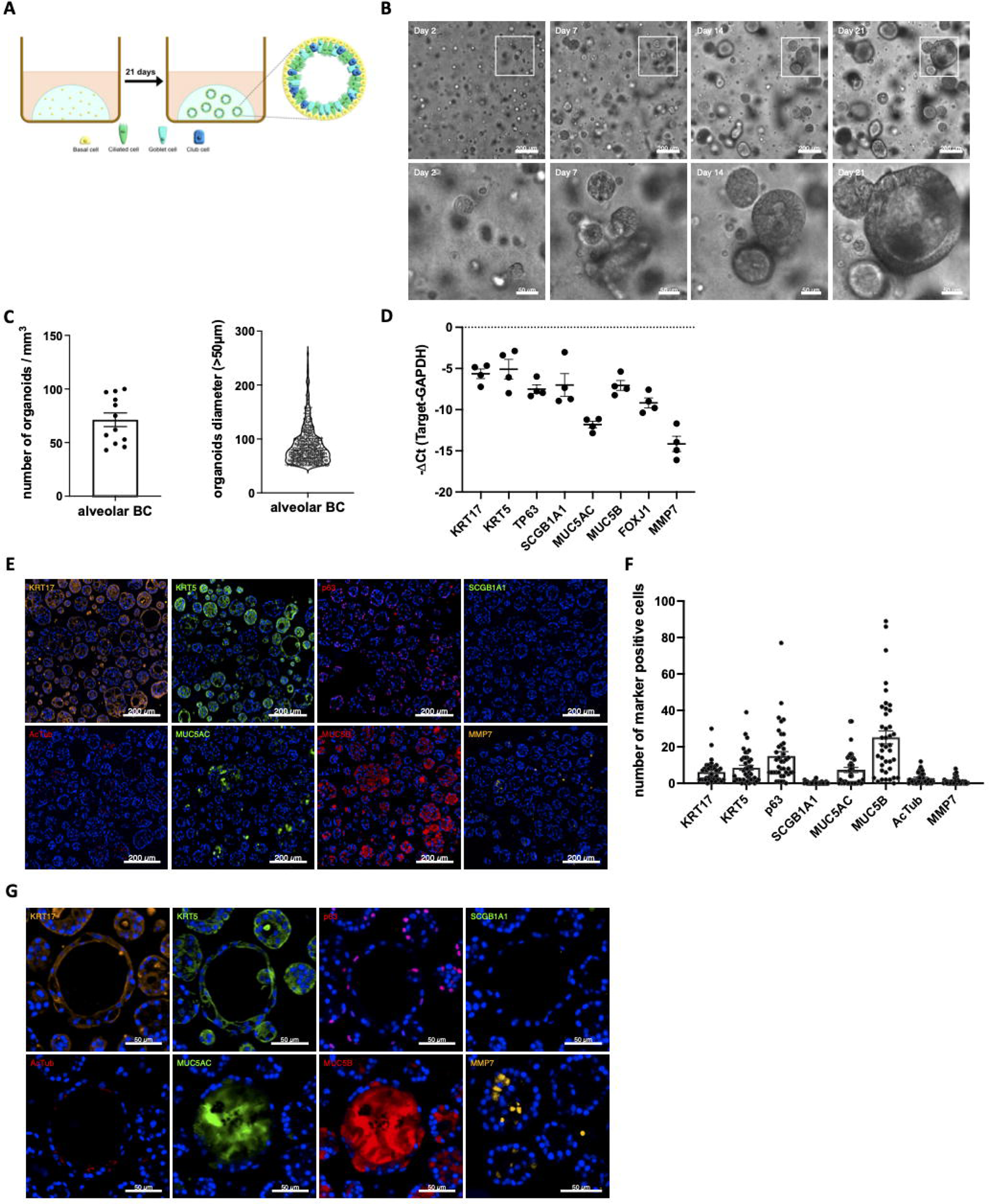
Alveolar basal cells form organoids and differentiate to secretory- and ciliated epithelial cells. Alveolar BC were resuspended in Matrigel and cultured for 21 days as illustrated (A). Experiments presented in this figure were all performed in alveolar BC from 4 IPF patients (n-number). Representative phase contrast images of organoid formation at day 2, 7, 14, and 21 (B). Graphs, showing the number of organoids/mm^3^ (mean ± SEM) and the distribution of their diameters >50μm (violin plot) of three selected regions (1mm^3^) per IPF patient (n=4) after 20 days (C). RNA expression levels (-ΔCt) of KRT17, KRT5, TP63, SCGB1A1, MUC5AC, MUC5B, FOXJ1, or MMP7 in organoids after 21 days. Data are presented as mean ± SEM (D). Representative ICC/IF images of organoids incubated with antibodies detecting KRT17, KRT5, p63, SCGB1A1, AcTub, MUC5AC, MUC5B, or MMP (E), and their quantification (mean ± SEM). Dots within the bar charts represent individual data points generated by analysis of 10 organoids per IPF patient (n=4) (F). Representative ICC/IF images of organoids incubated with antibodies detecting KRT17, KRT5, p63, SCGB1A1, AcTub, MUC5AC, MUC5B, or MMP7 (G). Nuclei were counter-stained with DAPI (G).

### Human alveolar BC engraft and differentiate in lung tissue of bleomycin-challenged mice

Mice were intratracheally challenged with a low dose of bleomycin for three days and were subsequently intratracheally injected with human alveolar BC or human alveolar BC transduced with a luciferase and GFP encoding vector. Mice were sacrificed after 21 days (Figure 6 A). Luciferase expression in mice injected with cells carrying a luciferase encoding vector, increased over time, suggesting pulmonary engraftment and proliferation of human alveolar BC (Figure 6 B, C). H&E stainings showed mild fibrotic changes in lung tissue of bleomycin-challenged mice (Figure 6 D). In lung tissue of mice injected with human alveolar BC, we observed extensive accumulation of cell with squamous morphology (Figure 6 D). Human BC invaded the alveolar space and partly destroyed alveolar septa (Figure 6 E). Cells stained positive for the human-specific protein HNA, confirming their human origin (Figure 6 F). IF stainings for basal-(KRT5, KRT17), secretory-(MUC5AC, MUC5B, SCGB1A1), or ciliated (AcTub) epithelial cell markers revealed that the human BC maintain their basal cell identity and differentiated towards secretory MUC5AC+, MUC5B+, SCGB1A1+ epithelial cells (Figure 6 F).

**Figure 6:**
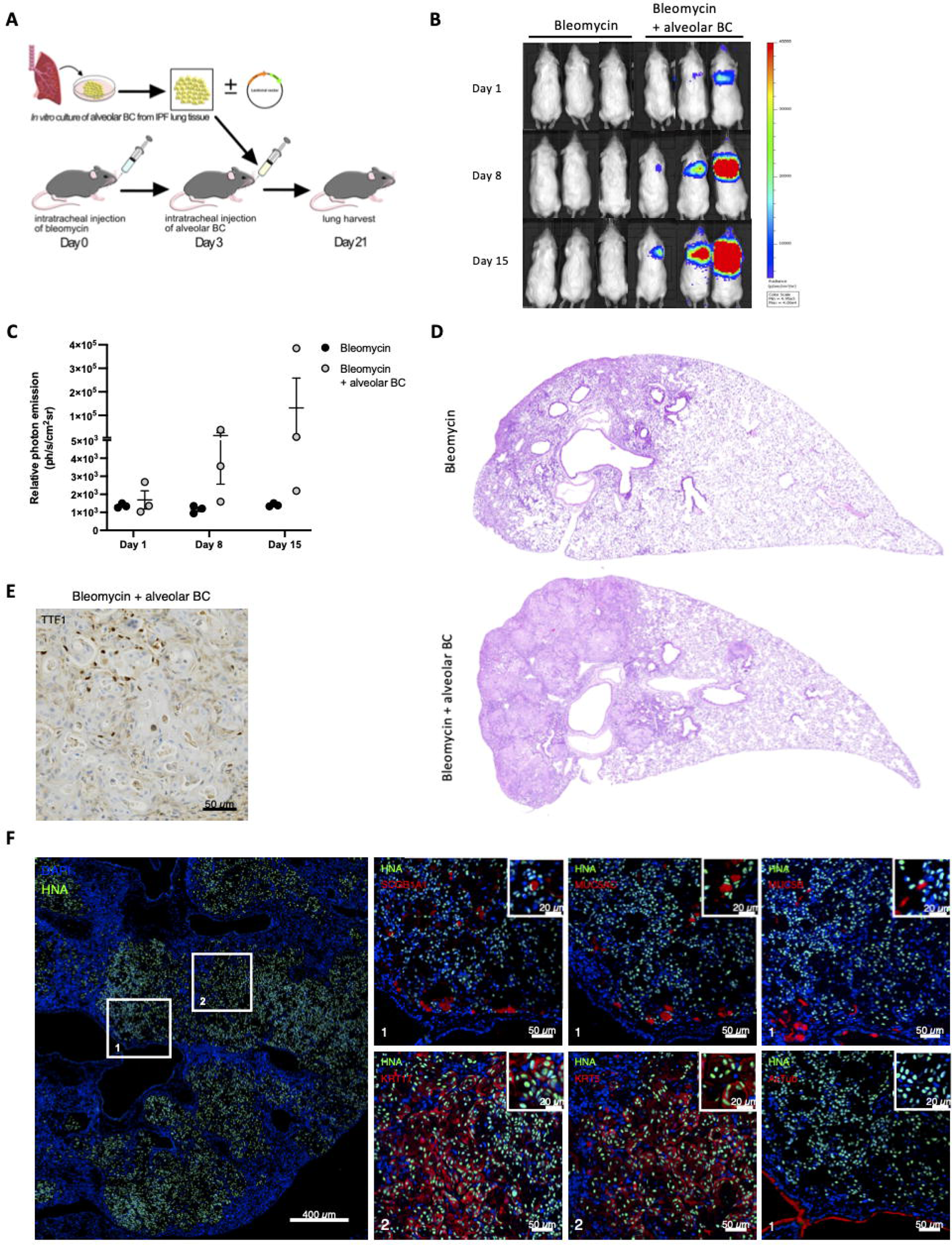
Human alveolar basal cells form HC-like structures in bleomycin-challenged mouse lungs. NRG mice were treated with bleomycin (n=3), or bleomycin + human alveolar BC carrying a luciferase encoding vector (n=3), or bleomycin + human alveolar BC (n=3) as illustrated (A). Luciferase expression in mice treated with bleomycin or bleomycin + human alveolar BC carrying a luciferase encoding vector (B) and the corresponding bioluminescence measurements (C) at day 1, 8, and 15. Representative images of H&E stainings in mice treated with bleomycin or bleomycin + human alveolar BC (D). Representative IHC image of TTF-1 in mice treated with bleomycin + human alveolar BC (E). Representative IHC/IF images of mouse lung tissue (mice treated with bleomycin + human alveolar BC) incubated with antibodies detecting human-specific HNA with or without SCGB1A1, MUC5AC, MUC5B, KRT17, KRT5, or AcTub. Nuclei were counter-stained with DAPI. White squares (1, and 2) indicate regions imaged at higher magnification (F).

## Discussion

In IPF, the alveolar lung parenchyma is replaced by dense fibrotic tissue and HC that are lined with a bronchiolar-like epithelium (8). In line with this, we here reported reduced numbers of SP-C+ AT2 cells, accompanied with the appearance of ectopic airway epithelial cells including basal-, ciliated-, and secretory epithelial cells within peripheral IPF lung tissue. Our findings are in agreement with several recent studies, showing dramatic changes in epithelial cell subtypes and the emergence of airway epithelial- and fibrosis-specific intermediate cell types, which mainly localize within HC in the IPF lung (4, 5, 9, 13–18). Although, the appearance of airway epithelial cells in the peripheral IPF lung is a widely accepted phenomenon, the cells origin remains elusive: Migration of airway epithelial cells into the alveolar space after injury (19), trans-differentiation of resident AT2 cells into alveolar BC (14, 15) or the occurrence of both mechanisms in parallel have been proposed (17). In addition, the cells functional role in disease development and progression is largely under-investigated. In animal models, basal-like cells helped to regenerate the injured alveolar epithelium (20, 21), whereas alveolar BC in the lung of IPF patients were associated with increased mortality and pathological HC formation (4, 22). By using IPF patient-derived alveolar BC, we here established *in vitro* and *in vivo* models that closely resemble HC formation in IPF. The use of such disease models will greatly contribute to a better understanding of the role and function of alveolar BC in IPF.

We previously described the fibrosis-enriched outgrowth and *the vitro* culture of alveolar BC (6, 10, 11). Under the previously described culture conditions, alveolar BC underwent growth arrest after a short culture period and spontaneously differentiated towards secretory- or aberrant basaloid-like cells, making long-term expansion and experimental use of the cells difficult (6, 11). We previously cultured alveolar BC in DMEM growth medium (6, 11), which contains high concentrations of calcium and likely lacks factors essential for the growth of epithelial cells. High calcium concentrations were shown to inhibit growth and to promote differentiation of epithelial cells (23, 24), likely explaining the cells insufficient proliferation and spontaneous differentiation in DMEM growth medium (6, 11). In this study, we replaced DMEM growth medium with the commercially available Cnt-PR-A medium, which was specifically developed to support growth and to inhibit differentiation of primary epithelial cells. Indeed, when cultured in Cnt-PR-A, alveolar BC expressed high levels of the canonical basal cell markers KRT5, KRT17, KRT14, and p63 and showed the capacity for robust proliferation and wound healing. Furthermore, spontaneous differentiation of alveolar BC was not observed under the optimized culture conditions and we were able to cryopreserve and expand the cells for several passages without loss of BC-specific markers.

We next aimed to establish *in vitro* models that closely resemble HC in the IPF lung, by using IPF-patient-derived alveolar BC. IF stainings of peripheral IPF lung tissue, confirmed the presence of KRT5+/KRT17+ basal-, AcTub+ ciliated- and SCGB1A1+, MUC5B+ and MUC5AC+ secretory epithelial cells within HC in the IPF lung. In accordance with other studies (5, 9), KRT5+/KRT17+, and MUC5B+ cells were the most prevalent cell types, whereas MUC5AC+ cells were detected only in small numbers. It was previously shown that airway BC differentiate to ciliated- and secretory epithelial cells when cultured on an ALI platform (25), whereas submerged culture of the cells prevented ciliary differentiation (26). We therefore cultured IPF-derived alveolar BC on an ALI platform, which induced their differentiation to ciliated- and secretory epithelial cells. In addition, we embedded alveolar BC into Matrigel in which they formed 3D organoids. Similar to our observations in ALI culture, alveolar BC-derived organoids were composed of basal-, ciliated-, and secretory epithelial cells. The cellular composition of alveolar BC-derived organoids closely resembled that of HC *in vivo*, with KRT5+/KRT17+ basal- and MUC5B+ cells being the most prevalent cell types.

Next, we examined alveolar BC behaviour after instillation into bleomycin-challenged mice. It was previously shown that airway-derived BC instilled in bleomycin-challenged mice, form HC-like structures within mouse lung tissue (22). Using the same animal model and protocol (22), we here show that IPF-patient derived alveolar BC engraft and proliferate in the lungs of bleomycin-challenged mice. In our study, the human BC-driven HC formation within mouse lungs was less pronounced. However, we showed that human alveolar BC differentiate to secretory epithelial cells within mouse tissue, suggesting a beginning formation of HC. The here and previously described humanized mouse model (22) therefore serves as a valuable model to study basal cell-driven HC formation *in vivo*.

In summary, the here described *in vitro* and *in vivo* models represent powerful tools to study HC formation in IPF. Although, our models may not fully reflect the *in vivo* situation of IPF HC, they appear superior to previously used models (22, 25) as BC, used in this study, 1) are IPF patient-derived and 2) were isolated from fibrotic IPF parenchyma, and not from the airways. Using cells from patient-derived tissue and from the region of interest seems important as it was shown that cells maintain disease- and region-specific characteristics when cultured *in vitro* (27, 28). Although our current study does not provide new insights into the origin or function of alveolar BC in IPF, the use of the here presented models in future studies will greatly enhance our knowledge about the role of alveolar BC in HC formation and its potential pharmacological inhibition in IPF.

## Experimental procedures

### Basal cell culture

Alveolar BC were cultured from lung-explants derived from IPF patients undergoing lung transplantation as previously described (6, 11). Briefly, lung tissue was cut into small pieces and placed into cell culture-treated plastic dishes containing the commercially available epithelial cell growth medium Cnt-PR-A. After 5-7 days the tissue pieces were removed and cells trypsinized and expanded. All experiments in this study were performed with cells between passage 1-2, except for the experiment evaluating the cell marker expression in cells between passage 0-6. For cryopreservation, confluent alveolar BC were trypsinized, re-suspended in cryo-medium (Cnt-PR-A 40%, fetal bovine serum 50%, DMSO 10%), stored in a -80° C freezer for up to one week and then transferred to liquid nitrogen tank for long-term storage. IPF was diagnosed based on ATS/ERS guidelines (1, 29). The local ethical committee of the University Hospital, Basel, Switzerland (EKBB05/06) approved the culture of human primary lung cells. Patient characteristics can be found in table S1. All materials used in this study are listed in table S2.

### TaqMan RT-PCR, Immunoblotting, Immunohistochemistry, immunocytochemistry, and Immunofluorescence

TaqMan® RT-PCR, immunoblotting, immunocytochemistry (ICC)/immunofluorescence (IF), immunohistochemistry (IHC)/IF or hematoxylin and eosin (H&E) stainings were performed as previously described (6, 11, 30). TaqMan PCR results are expressed as -ΔCt values (-(target-GAPDH)). Missing -ΔCt values (because the target genes was expressed at too low level to be detected after 40 cycles) were arbitrarily set to -20. Details for primers and antibodies are listed in table S2.

### Image acquisition and image processing

Lung tissue slides from non-fibrotic control- and IPF patients (see patient characteristics in table S1) were acquired as followed: Images of lung tissue were captured with Photometrics Prime 95B camera using a Nikon Plan Apo 20x objective (0.75 NA) at Nikon Ti2-E widefield microscope and stitched together to one large image (2555 x 2555 µm, pixel size 0.55 µm / pixel) using Nikon NIS Elements 4.21 software. Fluorophores (Alexa488, Alexa555, Alexa647) with 488 nm, 555nm, 647nm lasers respectively and a band pass emission filter (LED-DA/FI/TR/Cy5/Cy/-5x-A) were used. A total number of six large images of each IPF- or non-fibrotic control patient-derived lung tissue were analyzed. All acquired images were stored on the client-server platform Omero (Omero.insight 5.5.17 and Omero.web 5.9.1) for managing microscope images.

IF images were analyzed with Qupath version 0.3.4 (an open-source software for bioimage analysis) (31) using a script for each target protein including nuclei segmentation, pixel- and object classification. StarDist 2D (model: *dsb2018_heavy_augment.pb*), a deep-learning based-method of nucleus detection were used for IF images (32). All parameters used for cell detection in Qupath are listed in table 1. Following nuclei segmentation, the data were further classified for each marker (cytosol, nucleus or membrane depending on the target labeling) and applied to all datasets. For each marker, thresholds were set visually and mean intensity was assessed to quantify positive cells in lung tissue. Data is presented as mean ± SEM.

**Table 1:** Parameters used in Qupath script.

| Parameters | Values |
| --- | --- |
| Probability (detection) threshold | 0.5 |
| Selected detection channel | DAPI |
| Percentile normalization | 1, 99 |
| Resolution for detection (pixel size) | 0.275 |
| Cell expansion | 1.0 |
| Cell constrains scale | 1.5 |
| Probability | True |
| **Intensity threshold parameters (localization)** |  |
| SP-C (cytosol) | 800 |
| KRT17 (cytosol) | 400 |
| KRT5 (cytosol) | 400 |
| KRT14 (cytosol) | 700 |
| p63 (nucleus) | 1000 |
| AcTub (membrane) | 400 |
| SCGB1A1 (cytosol) | 400 |
| MUC5AC (cytosol) | 300 |
| MUC5B (cytosol) | 300 |
| MMP7 (cytosol) | 600 |

### Air liquid interface culture

For air liquid interface culture (ALI), alveolar BC were seeded on 12 mm transwell inserts (apical chamber) in a 12-well plate (basal chamber) and grown to confluence by adding Cnt-PR-A to the basal and apical chamber. To initiate differentiation, Cnt-PR-A medium was aspirated from both chambers. PneumaCult-ALI maintenance medium was prepared according to the manufacturer’s instructions and added to the basal chamber only. Medium in the basal chamber was changed every 2-3 days for a total of 23 days. The cells on the transwell membranes were either lysed for RNA expression analysis or fixed in 4 % paraformaldehyde (PFA) and analyzed by ICC/IF. Alternatively, the cells were fixed in 4 % PFA for 15 min at RT, embedded in Tissue-Tek^®^ O.C.T. compound after 30 % and 15 % sucrose incubation and snap frozen in 2-methylbutan with liquid nitrogen. Frozen sections were cut into 7 µm on a cryostat and analyzed by ICC/IF.

### 3D organoid culture

Cultured alveolar BC were trypsinized and resuspended in 40 μl ice-cold Matrigel at a density of 500 cells/μl. Droplets were placed in a pre-warmed 24-well suspension plate and incubated for 20 min (37°C, 5 % CO_2_). Differentiation media formulations were slightly modified from a previously described protocol (33): Basic differentiation media (bDM) consists of IMDM-Ham’s F12 (3:1 vol/vol) with B 27 (with retinoic acid) supplement (1 %), N 2 supplement (0.5 %), BSA (0.05 %), antibiotic-antimycotic 1x, glutamax 1x, ascorbic acid (50 μg/ml) and monothioglycerol (4.5×10^-4^ M). Complete differentiation medium (cDM) consists of bDM supplemented with FGF-2 (250 ng/ml), FGF-10 (100 ng/ml), EGF (5 ng/ml), dexamethasone (50 nM), 8-Bromo-cAMP (0.1 mM) IBMX (0.1 mM), and A83-01 (TGF-ß inhibitor, 1 μM). For the first 7 days, a mixture of epithelial cell growth medium (Cnt-PR-A) and cDM (1:1) with ROCK inhibitor (Y-27632 dihydrochloride, 10 μM) was added to the wells. The medium was then changed to cDM and incubated for another 14 days. cDM was exchanged once after 7 days. On day 20 droplets with organoids were imaged at 10x with a z-stack (every 50 µm) and stitched together by Nikon Ti2-E widefield microscope. The diameter and number of organoids per mm^3^ with a diameter >50 μm as well as the percentage of organoids with polarized lumen was determined by using Fiji software version 2.9.0 (34). On day 21 the Matrigel was digested with Dispase II (2mg/ml) for 20min at 37°C and organoids were fixed in 4 % PFA for 30 min, embedded in pre-heated histogel, polymerized for 1 hour at RT and stored in 50 % ethanol for up to 1 week. Organoids were then dehydrated using ethanol series (50%, 70%, 95%, 100%) followed by Xylene incubation and then embedded in paraffin. Paraffin blocks were cut on a microtome into 4 µm sections and analysed by ICC/IF. For ICC/IF image quantification, a selection of organoids (n = 10) across each IPF patient (n=4) was captured at 40x using a Nikon Ti2-E widefield microscope. Automated counting of single-color images (each channel separately) for target proteins of each organoid was performed in FIJI. Alternatively, organoids were lysed for RNA expression analysis.

### Proliferation assay

Alveolar BC were seeded into two wells of a 24-well plate. When the cells reached about 70% confluence, one of the wells was stained with live cell stain NucBlue, imaged at 4x using a Nikon Ti2-E widefield microscope and the cells counted by using the general analysis NIS software version 4.2.1 (time point 0). The same procedure was repeated with the second well after 48 hours.

### Epithelial wound repair assay

Alveolar BC were seeded into 12-well culture plates and grown to confluence. The cell layer was mechanically wounded by using a pipette tip. Wound closure was monitored by using a Nikon Ti2 2.3.PO widefield time lapse microscope and calculated by using a Fiji plugin wound healing size tool (35).

### Intratracheal administration of human alveolar BC into bleomycin-challenged mice

All mouse procedures were conducted at the Hannover Medical School, Germany in accordance with the German law for animal protection and the European Directive 2010/63/EU and were approved by the Lower Saxony State Office for Consumer Protection and Food Safety in Oldenburg/Germany (LAVES); AZ: 33.12-42502-04-15/1896 and AZ: 33.19-42502-04-15/2017), AZ:33.12-42502-04-17/2612). Human alveolar BC were cultured as described earlier and some of the cells were transduced with a lentiviral vector (kindly provided by Dr. Axel Schambach, Hannover Medical School) for firefly-luciferase and enhanced green fluorescence protein (eGFP) as previously described (22). A total of nine NRG mice received bleomycin at a dose of 1.2 mg/kg intratracheally at day 0. Three days later, three mice were intratracheally injected with human alveolar BC, and three mice with vector-transduced human alveolar BC (in both conditions: 0.3 × 10^5^ cells per mouse). For *in vivo* imaging of firefly-luciferase activity in vector-transduced human alveolar BC, the respective mice were injected subcutaneously with XenoLight D-Luciferin-K+ Salt Bioluminescent Substrate (150 mg/kg) on day 1, 8 and 15. Bioluminescence was subsequently measured by an IVIS Lumina II and data analyzed using LivingImage 4.5 as previously described (22). At day 21 mouse lungs were harvested and embedded in paraffin for IHC/IF analysis.

### Statistics

Experiments were performed in alveolar BC derived from multiple different IPF patients as indicated by the n-numbers for each individual experiment. Statistical analysis was performed using GraphPad Prism software version 9.1.1. Paired or unpaired t-test was applied to determine p values, and the data are presented as mean ± SEM. All p values < 0.05 were considered as statistically significant.

## Acknowledgments

The authors acknowledge the microscopy core facility at the Department of Biomedicine in Basel, Switzerland for their support and assistance in ICC/IF, IHC/IF image acquisition and quantification.

Dr. Axel Schambach from Hannover Medical School kindly provided the lentiviral vector.

This project was supported by a project grant (310030_192536) by the Swiss National Research Foundation.

## Author Contributions

**SB**: Cell culture of human primary lung cells, ALI and organoid processing, ICC/IF, IHC/IF, image acquisition and image quantification in cultured cells, human IPF- and mouse tissue slides, immunoblotting, PCR, proliferation assay, scratch assay, data analysis and interpretation, revision and editing of the manuscript; **PK:** Conception and design of the study, cell culture of human primary lung cells, PCR, data analysis and interpretation, writing the manuscript; **NA:** bleomycin-mouse model, data analysis and interpretation, revision and editing of the manuscript for important intellectual content; **LP:** bleomycin-mouse model, data analysis and interpretation, revision and editing of the manuscript for important intellectual content; **SS:** Paraffin embedding, cutting, and H&E stainings of non-fibrotic control- and IPF lung tissue, interpretation of stainings performed on human IPF- and mouse lung tissue slides, revision and editing of the manuscript for important intellectual content; **LK:** Provided samples of IPF lung explants, revision and editing of the manuscript. **DJ:** Provided samples of IPF lung explants, revision and editing of the manuscript. **MK:** Provided samples of IPF lung explants, revision and editing of the manuscript; **AP:** bleomycin-mouse model, data analysis and interpretation, revision and editing of the manuscript for important intellectual content; **KEH:** Conception and design of the study, data analysis and interpretation, revision and editing of the manuscript for important intellectual content.

## Supplemental information

Patient characteristics in table S1. All materials used in this study are listed in table S2.

