## Supplement Table 1, 2 for "The use of cultured human alveolar basal cells to mimic honeycomb formation in idiopathic pulmonary fibrosis"

**Table S1: Patient Characteristics**

| Patient ID | Sex | Age (years) | Clinical Diagnosis |
|---|---|---|---|
| 01 | Male | 34 | fibrosis - UIP |
| 02 | Male | 60 | IPF |
| 03 | Male | 57 | IPF |
| 04 | Male | 49 | fibrosis - UIP |
| 05 | Female | 62 | IPF |
| 06 | Male | 54 | IPF |
| 07 | Male | 56 | IPF |
| 08 | Male | 60 | IPF |
| 09 | Mal | 54 | IPF |
| 10 | Female | 61 | COPD/control |
| 11 | Female | 74 | Cancer/control |
| 12 | Female | 70 | Cancer/control |
| 13 | Female | 61 | Cancer/control |

*Abbreviations used:* IPF, idiopathic pulmonary fibrosis; UIP, usual interstitial pneumonia.

**Table S2: Materials used in this study**

| Product | Company | Catalog Number | Country |
|---|---|---|---|
| Cnt-PR-A | CELLnTEC | - | Bern, Switzerland |
| 0.05% Trypsin-EDTA | Thermo Fisher Scientific | 25300-054 | Waltham, MA, USA |
| PneumaCult$^{TM}$-ALI medium | Stemcell Technology | 05001 | Cambridge, UK |
| Costar® 12 mm Transwell®, 0.4 μm pore polyester membrane inserts | Corning | 38024 | NY, USA |
| Heparin Solution | Stemcell Technology | 07980 | Cambridge, UK |
| Hydrocortisone stock solution | Stemcell Technology | 07925 | Cambridge, UK |
| Cultrex$^{TM}$ reduced growth factor basement membrane extract, type 2 | RnD Systems | 3533-005-02 | Abingdon, UK |
| IMDM | Thermo Fisher Scientific | 12440053 | Waltham, MA, USA |
| F12 nutrient mix, Hams | Thermo Fisher Scientific | 11765054 | Waltham, MA, USA |
| B 27 supplement | Thermo Fisher Scientific | 17504044 | Waltham, MA, USA |
| N 2 supplement | Thermo Fisher Scientific | 17502048 | Waltham, MA, USA |
| BSA (7.5% stock) | Thermo Fisher Scientific | 15260037 | Waltham, MA, USA |
| Glutamax 100X | Thermo Fisher Scientific | 35050061 | Waltham, MA, USA |
| Ascorbic Acid (50 mg/ml stock) | Sigma Aldrich | A4544 | Buchs, Switzerland |
| Antibiotic-antimycotic (100x) | Thermo Fisher Scientific | 15240-062 | Waltham, MA, USA |
| Monothioglycerol | Sigma Aldrich | M6145 | Buchs, Switzerland |
| 8-Bromo-cAMP | Stemcell Technology | 73604 | Cambridge, UK |
| Recombinant human FGF-2 protein | RnD Systems | 233-FB-025 | Abingdon, UK |
| Recombinant human EGF protein | PreproTech | AF-100-15 | London, UK |
| Recombinant human FGF-10 protein | RnD Systems | 345-FG-025 | Abingdon, UK |
| A83-01 | RnD Systems | 2939/10 | Abingdon, UK |
| Y-27632 dihydrochloride | RnD Systems | 1254/10 | Abingdon, UK |
| Dexamethasone | Sigma Aldrich | D4902 | Buchs, Switzerland |
| IBMX | Stemcell Technology | 72762 | Cambridge, UK |

| | | | |
|---|---|---|---|
| Dispase II | Sigma Aldrich | D4693-1g | Buchs, Switzerland |
| Tissue-Tek® O.C.T. compound | Sakura | 4583 | Torrance, CA, USA |
| Mouse KRT17 antibody | Thermo Fisher Scientific | MA1-06325 | Waltham, MA, USA |
| Rabbit KRT17 antibody | Thermo Fisher Scientific | PA5-27949 | Waltham, MA, USA |
| Mouse KRT5 antibody | Thermo Fisher Scientific | MA5-12596 | Waltham, MA, USA |
| Rabbit proSP-C antibody | Merck Millipore | AB3786 | Darmstadt, Germany |
| Mouse anti-tubulin, acetylated antibody | Merck Millipore | T6793 | Darmstadt, Germany |
| Rabbit MMP-7 antibody | Thermo Fisher Scientific | PA5-87486 | Waltham, MA, USA |
| Rat SCGB1A1 antibody | RnD System | 394324 | Abingdon, UK |
| Goat p63 antibody | RnD System | AF1916 | Abingdon, UK |
| Rabbit p63 antibody | Abcam | ab124762 | Cambridge, MA, USA |
| Mouse N-cadherin (CDH2) antibody | Thermo Fisher Scientific | 33-3900 | Waltham, MA, USA |
| Rabbit E-cadherin (CDH1) antibody | Abcam | Ab40772 | Cambridge, MA, USA |
| Rabbit GAPDH antibody | Abcam | Ab128915 | Cambridge, MA, USA |
| Mouse MUC5AC antibody | Thermo Fisher Scientific | MA5-12178 | Waltham, MA, USA |
| Mouse KRT14 antibody | Thermo Fisher Scientific | MA5-11599 | Waltham, MA, USA |
| Rabbit HNA antibody | Abcam | Ab86129 | Cambridge, MA, USA |
| Hematoxyline Erythrosin (H&E) | RAL Diagnostics | - | Bordeaux, France |
| TTF-1 antibody | Ventana | 790-4398 | Tucson, AZ, USA |
| Rabbit MUC5B antibody | Merck Millipore | HPA008246 | Darmstadt, Germany |
| Alexa 647 Goat anti-mouse | Thermo Fisher Scientific | A21235 | Waltham, MA, USA |
| Alexa 488 Donkey anti-mouse | Thermo Fisher Scientific | A21202 | Waltham, MA, USA |
| Alexa 488 Donkey anti-rat | Thermo Fisher Scientific | A21208 | Waltham, MA, USA |
| Alexa 488 Goat anti-rabbit | Thermo Fisher Scientific | A11008 | Waltham, MA, USA |
| Alexa 555 Goat anti-rabbit | Thermo Fisher Scientific | A21428 | Waltham, MA, USA |
| Alexa 594 Donkey anti-goat | Thermo Fisher Scientific | A11058 | Waltham, MA, USA |
| Alexa 647 Goat anti-rat | Thermo Fisher Scientific | A21247 | Waltham, MA, USA |

| | | | |
|---|---|---|---|
| DAPI | Thermo Fisher Scientific | 62248 | Waltham, MA, USA |
| Donkey anti-goat Peroxidase antibody | Sigma Aldrich | SAB3700284 | Buchs, Switzerland |
| Goat anti-rabbit Peroxidase antibody | Sigma Aldrich | A9169 | Buchs, Switzerland |
| Goat anti-mouse Peroxidase antibody | Sigma Aldrich | A9917 | Buchs, Switzerland |
| Nikon Ti2-E widefield microscope | Nikon | - | - |
| Nikon Ti2 2.3.PO widefield microscope | Nikon | - | - |
| Nikon Ni – Slide scanner | Nikon | - | - |
| KRT5 TaqMan® Gene Expression Assay | Thermo Fisher Scientific | Hs00361185_m1 | Waltham, MA, USA |
| KRT17 TaqMan® Gene Expression Assay | Thermo Fisher Scientific | Hs00356958_m1 | Waltham, MA, USA |
| GAPDH TaqMan® Gene Expression Assay | Thermo Fisher Scientific | Hs03929097_g1 | Waltham, MA, USA |
| TP63 TaqMan® Gene Expression Assay | Thermo Fisher Scientific | Hs00978340_m1 | Waltham, MA, USA |
| KRT14 TaqMan® Gene Expression Assay | Thermo Fisher Scientific | Hs00265033_m1 | Waltham, MA, USA |
| MUC5AC TaqMan® Gene Expression Assay | Thermo Fisher Scientific | Hs01365616_m1 | Waltham, MA, USA |
| FOXJ1 TaqMan® Gene Expression Assay | Thermo Fisher Scientific | Hs00230964_m1 | Waltham, MA, USA |
| SCGB1A1 TaqMan® Gene Expression Assay | Thermo Fisher Scientific | Hs00171092_m1 | Waltham, MA, USA |
| CDH1 TaqMan® Gene Expression Assay | Thermo Fisher Scientific | Hs01023894 | Waltham, MA, USA |
| CDH2 TaqMan® Gene Expression Assay | Thermo Fisher Scientific | Hs00983056_m1 | Waltham, MA, USA |
| MUC5B TaqMan® Gene Expression Assay | Thermo Fisher Scientific | Hs06629268_s1 | Waltham, MA, USA |
| TaqManTM Universal PCR Master Mix, no AmpEraseTM UNG | Thermo Fisher Scientific | 4324018 | Waltham, MA, USA |
| Quick-RNA MiniPrep Kit | ZymoResearch | R1050 | Orange, CA, USA |
| XenoLight D-Luciferin-K+ Salt Bioluminescent Substrate | PerkinElmer | 122799 | USA |
| IVIS Lumina II | PerkinElmer | - | - |
| LivingImage 4.5 | PerkinElmer | - | - |

| | | | |
|---|---|---|---|
| NucBlue Live Cell Stain Ready Probes | Thermo Fisher Scientific | R37605 | Waltham, MA, USA |
